# Tmem117 in AVP neurons regulates the counterregulatory response to hypoglycemia

**DOI:** 10.1101/2022.10.21.513159

**Authors:** Sevasti Gaspari, Gwenaël Labouèbe, Alexandre Picard, Xavier Berney, Ana Rodriguez Sanchez-Archidona, Bernard Thorens

## Abstract

The counterregulatory response to hypoglycemia (CRR), which ensures a sufficient glucose supply to the brain, is an essential survival function. It is orchestrated by incompletely characterized glucose-sensing neurons, which trigger a coordinated autonomous and hormonal response that restores normoglycemia. Here, we investigated the role of hypothalamic *Tmem117*, identified in a genetic screen as a regulator of CRR. We show that *Tmem117* is expressed in vasopressin magnocellular neurons of the hypothalamus. *Tmem117* inactivation in these neurons increases hypoglycemia-induced vasopressin secretion leading to higher glucagon secretion, an estrus cycle phase-dependent effect in female mice. *Ex vivo* electrophysiological analysis, in-situ hybridization and *in vivo* calcium imaging reveal that *Tmem117* inactivation does not affect the glucose-sensing properties of vasopressin neurons but increases ER-stress, ROS production and intracellular calcium levels accompanied by increased AVP production and secretion. Thus, *Tmem117* in vasopressin neurons is a physiological regulator of glucagon secretion and highlight the role of these neurons in the coordinated response to hypoglycemia.

## Introduction

The brain relies almost exclusively on glucose for metabolic energy production. Therefore, maintaining blood glucose levels no lower than ∼5 mM is essential for survival (Marty, Dallaporta and Thorens, 2007). When glycemic levels drop below this threshold, a brain-orchestrated neuroendocrine reflex triggers a counterregulatory response (CRR). This is characterized by the secretion of multiple hormones that act on peripheral target organs to stimulate endogenous glucose production, suppress insulin secretion and minimize glucose utilization to restore normoglycemia (Tesfaye and Seaquist, 2010). A key aspect of the CRR is the induction of glucagon (GCG) secretion from pancreatic alpha cells, which triggers enhanced glucose production from the liver (Ramnanan *et al*., 2011; Thorens, 2022). Patients with type 1 or advanced type 2 diabetes display loss of hypoglycemia-induced GCG secretion (Cryer, 2012; Siafarikas *et al*., 2012; Bisgaard Bengtsen and Møller, 2021), but the underlying mechanism is poorly understood. In addition to deregulated alpha-cell autonomous, and intra-islet paracrine interactions (Gaisano, MacDonald and Vranic, 2012), evidence suggests that defects in the central nervous system (CNS) control of GCG secretion contribute to this blunted response (Beall, Ashford and McCrimmon, 2012; Stanley, Moheet and Seaquist, 2019).

In the CNS, changes in glucose concentrations are detected by neurons located in several regions but have been studied most extensively in the hypothalamus and brainstem. Several hypothalamic areas are equipped with glucose-sensing neurons, including the ventromedial (VMH), dorsomedial (DMH), lateral, arcuate (ARC), paraventricular (PVN) and supraoptic (SON) nuclei (Stanley, Moheet and Seaquist, 2019). Glucose responsive neurons are activated by either a rise (glucose excited / GE neurons) or a fall (glucose inhibited / GI neurons) in glucose levels. The response to hypoglycemia involves activation by GI neurons of both the parasympathetic and sympathetic branches of the autonomic nervous system, which stimulate GCG secretion and hepatic glucose production, the activation of the hypothalamus-pituitary-adrenal (HPA) axis to stimulate epinephrine and glucocorticoid secretion (Tesfaye and Seaquist, 2010), and th activation of hypothalamic AVP neurons to stimulate GCG secretion through the actions of AVP on pancreatic alpha cell AVP V1b receptors (Gao, Gérard and Henquin, 1992; Yibchok-anun *et al*., 2004; Kim *et al*., 2021).

Multiple studies have provided evidence in support for the role of hypothalamic and brainstem glucose sensing neurons in CRR. For instance, cell-specific activation of VMH neuronal subpopulations expressing glucokinase (Gck) (Meek *et al*., 2016) or steroidogenic-factor 1 (Sf1) (Stanley *et al*., 2016) increases GCG secretion and glycemia, while their inhibition blocks CRR. Interestingly, genetic inactivation of *Gck* in Sf1 neurons impairs GCG secretion in a sex-dependent manner (Steinbusch *et al*., 2016). Glucose responsive neurons located in other nuclei have also been shown to control GCG secretion. This is the case of cholecystokinin-expressing neurons of the parabrachial nucleus that control VMH neurons (Garfield *et al*., 2014); of fibroblast growth factor 15-expressing neurons of the DMH (Picard *et al*., 2016; Picard *et al*., 2021); and of Glut2 GI neurons of the nucleus tractus solitarius, which activate vagal nerve firing to stimulate GCG secretion (Lamy *et al*., 2014).

To identify in an unbiased manner novel hypothalamic regulators of CRR induced by insulin-induced hypoglycemia, we performed a genetic screen using a panel of recombinant inbred BXD mouse lines, derived from the cross of C57Bl/6 and DBA/2 mice (Peirce *et al*., 2004). This screen (Picard *et al*., 2022) led to the identification of *Agpat5*, encoding a lipid biosynthesis enzyme, and the characterization of its essential role in the activation by hypoglycemia of AgRP neurons and GCG secretion (Strembitska *et al*., 2022). We further identified *Tmem117*, located in a QTL on chromosome 15, as another candidate regulator of GCG secretion. *Tmem117* encodes an eight transmembrane-containing membrane protein (Bürgi *et al*., 2016) that has been reported to negatively control ER-stress and ROS production (Tamaki *et al*., 2017).

Here, we show that *Tmem117* is expressed in vasopressin (AVP) magnocellular neurons, that AVP neurons are in large part GI neurons, and that hypoglycemia induces copeptin (CPP; an AVP surrogate) and GCG secretion. Genetic inactivation of *Tmem117* in AVP neurons increases, in a sex-dependent manner, hypoglycemia-induced CPP and GCG secretion. *Tmem117* inactivation does not affect the glucose sensing properties of AVP neurons, but induces ER-stress and increases intracellular ROS and Ca^++^ levels as well as AVP mRNA expression.

## Results

### *Tmem117* is expressed in AVP magnocellular neurons

A genetic screen of a panel of 36 recombinant inbred BXD mice for hypoglycemia-induced GCG secretion identified a clinical quantitative trait locus (cQTL) on chromosome 8 and one on chromosome 15 (Picard *et al*., 2022). The candidate gene on chromosome 8 encodes the lipid-modifying enzyme Agpat5, which is required for hypoglycemia sensing by AgRP neurons, vagal nerve activation and GCG secretion (Strembitska *et al*., 2022). On chromosome 15 two genes showed strong negative correlation with the GCG trait, *Irak4* and *Tmem117* (Fig S1A-D). The role of *Irak4* in controlling hypothalamic Il-1β signaling and GCG secretion has been reported (Picard *et al*., 2022). Here, we performed an expression QTL analysis (eQTL) for the level of *Tmem117* mRNA in the hypothalamus of the BXD mouse lines. We identified an eQTL on chromosome 15 at the same position of the cQTL (Fig S1E). Thus, indicating that this genomic locus controls both GCG secretion and *Tmem117* expression.

**Figure S1.**
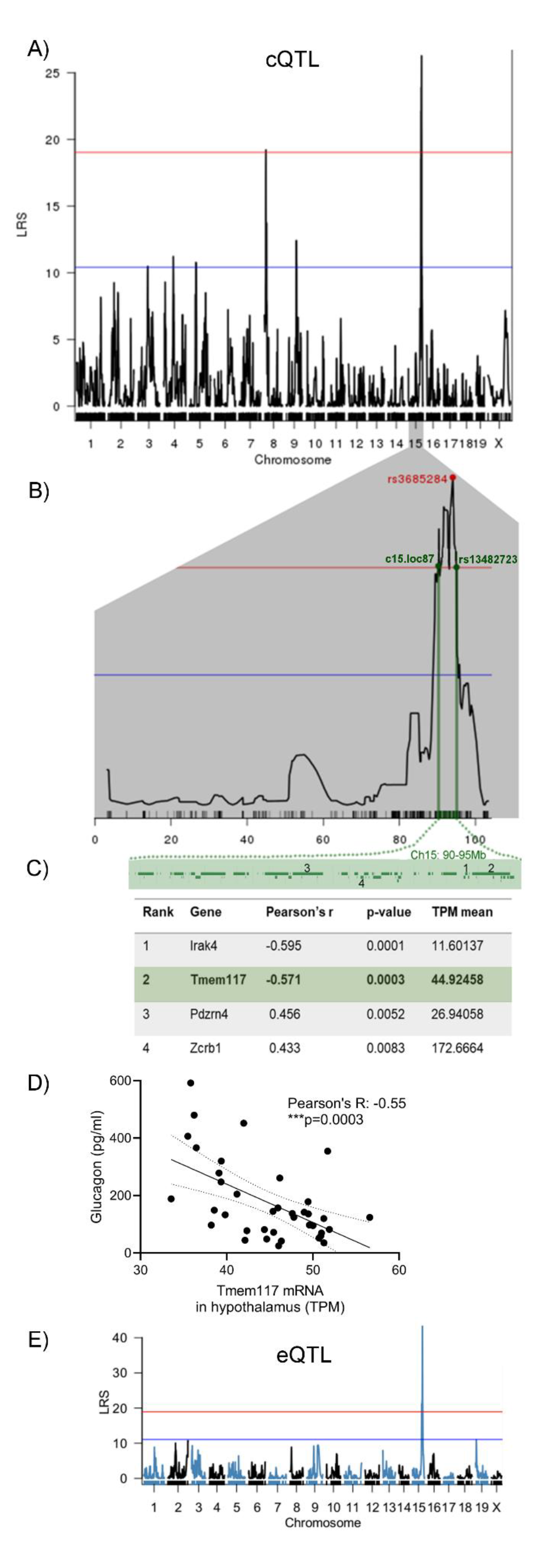
Hypothalamic *Tmem117* is a candidate gene for the control of insulin-induced glucagon secretion. (A) Clinical quantitative trait locus (cQTL) analysis revealed two genomic regions controlling insulin-induced GCG secretion, on chromosome 8 and 15. (B) The peak on chromosome 15, between markers c15.loc87 and rs13482723, contains 42 genes. (C) The genes showing the highest correlation between their hypothalamic mRNA expression and plasma GCG are *Irak4* and *Tmem117*. (D) Correlation between the level of *Tmem117* expression in the hypothalamus of the 36 BXD mouse lines and insulin-induced plasma GCG levels. (E) Expression quantitative trait locus (eQTL) analysis for *Tmem117* expression in the hypothalamus of the 36 BXD mouse strains. A cis eQTL was identified on chromosome 15 with a LRS of 43.3. Panels A-C are adapted from *Picard et al., 2022*. LRS: likelihood ratio statistic, TPM: transcripts per million.

To establish the sites of *Tmem117* expression in the hypothalamus we performed immunofluorescence microscopy analysis. We found strong immunolabeling for Tmem117 in the SON and PVN (Fig 1A) and in the posterior pituitary (Fig 1B). Co-staining for neuropeptides revealed that *Tmem117* was expressed in AVP magnocellular cells (Fig 1E-G). The specificity of the immunostaining was confirmed by the lack of signal in brains of mice with *Tmem117* gene inactivation in AVP neurons (*Tmem117^fl/fl^*;*AVP-IRES-Cre-D^tg/+^* mice) (Fig 1C,D). Quantification in three mice and two consecutive hypothalamic sections for each mouse (bregma −0.7 and −0.8) showed that ∼90 percent of AVP neurons in the PVN and ∼97 percent in the SON were Tmem117 positive (Fig 1H,I). Super-resolution microscopy revealed a punctuated intracellular distribution of Tmem117, which colocalized in part with AVP granules in both the soma and axons of magnocellular neurons (Fig 1J).

**Figure 1.**
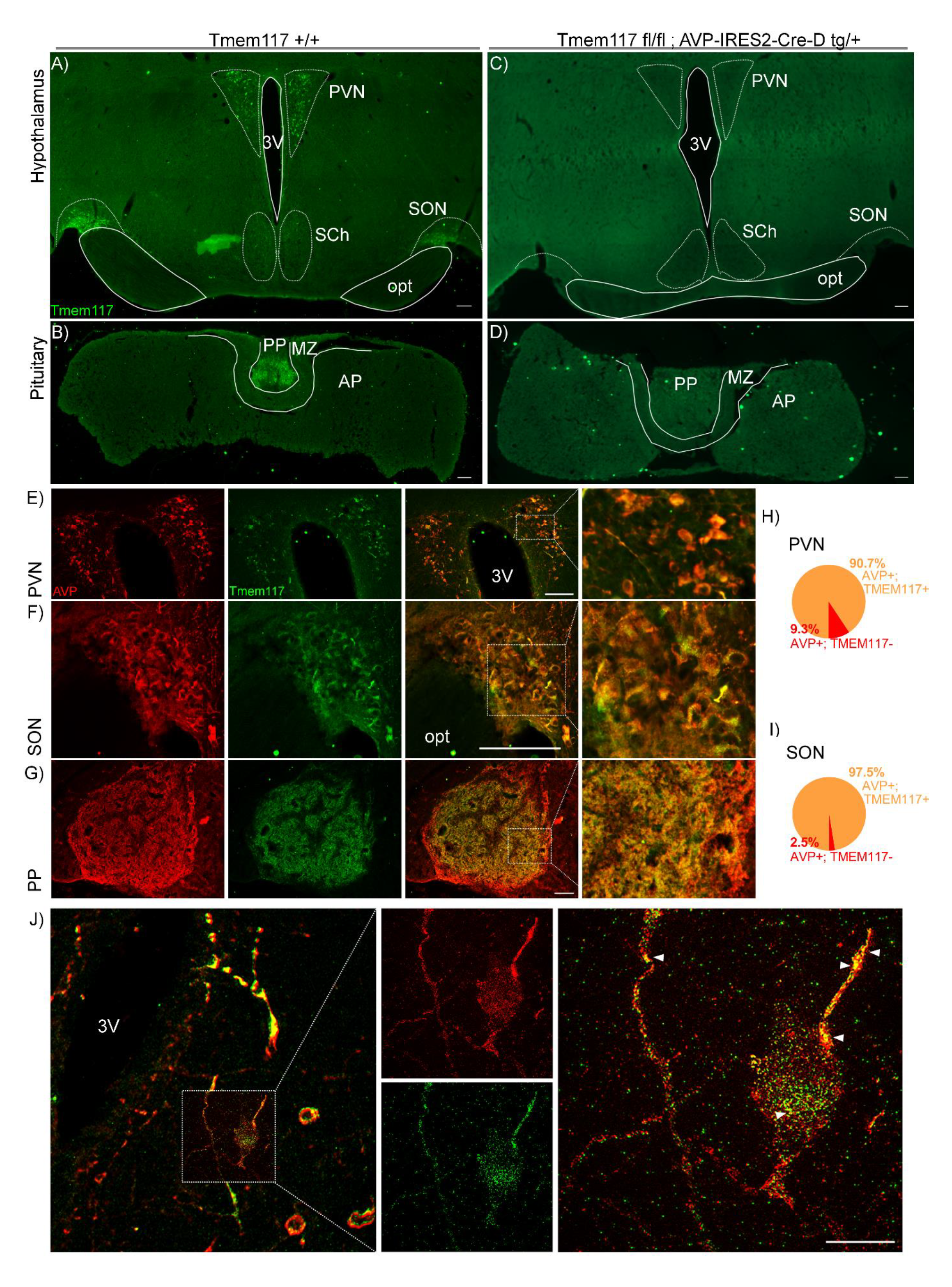
*Tmem117* is expressed in AVP magnocellular neurons. Immunofluorescence microscopy detection of Tmem117 in the mouse hypothalamus and pituitary: (A, B) Positive immunostaining was observed in the PVN, the SON and the posterior pituitary. (C, D) No immunostaining was detected in the hypothalamus and pituitary of mice with constitutive inactivation of *Tmem117* in AVP neurons. (E) Co-staining for Tmem117 and AVP in the PVN. (F) Co-staining for Tmem117 and AVP in the SON. (G) Co-staining for Tmem117 and AVP in the posterior pituitary. (H, I) Quantification of AVP^+^/Tmem117^+^ cells in the PVN and SON, respectively. (J) Higher resolution depiction of an AVP^+^/Tmem117^+^ neuron in the PVN captured with structured illumination microscopy. The white arrows highlight representative positions of signal co-localization. 3V: 3^rd^ ventricle, AP: Anterior pituitary, MZ: Medial zone, opt: optic tract, PP: Posterior pituitary, PVN: Paraventricular nucleus, SCh: Suprachiasmatic nucleus, SON: Supraoptic nucleus. Scale bar=100μm for panels A-G, =10μm for panel J.

### *Tmem117* inactivation in AVP neurons increases insulin-induced copeptin and glucagon secretion

To inactivate *Tmem117* specifically in AVP neurons, we generated *Tmem117^fl^* mice that allowed Cre-dependent excision of exon 3, which encodes the third transmembrane domain (amino acids 93-136) of the predicted structure of Tmem117 (Jumper *et al*., 2021; Varadi *et al*., 2022) (Fig S2A,B). To induce *Tmem117^fl^* recombination in AVP neurons, we used a viral-mediated approach for expression of Cre recombinase. This was preferred over the use of *AVP-IRES-Cre-D^tg/+^* mice because these display reduced endogenous AVP levels (Cheng, Fung and Cheng, 2019). We, thus, constructed an AAV plasmid containing the full-length AVP promoter placed upstream of a codon-improved Cre recombinase (iCre) sequence (Ponzio *et al*., 2012) (Fig S2C). The resulting plasmid was packaged in serotype 6 adeno-associated viruses (AAV6-AVP-iCre), which infect magnocellular neurons with high efficacy (Ponzio *et al*., 2012) and have a high retrograde transport capacity (Salegio *et al*., 2013). Injection of AAV6-AVP-iCre in the posterior pituitary of adult *Tmem117^fl/fl^* mice triggered efficient recombination in the PVN and SON as detected by PCR analysis of genomic DNA (Fig S2D) and by the expression of tdTomato when the AAV6-AVP-iCre was injected in the posterior pituitary of *Rosa26^tdTomato^* mice (Fig S2E).

**Figure S2.**
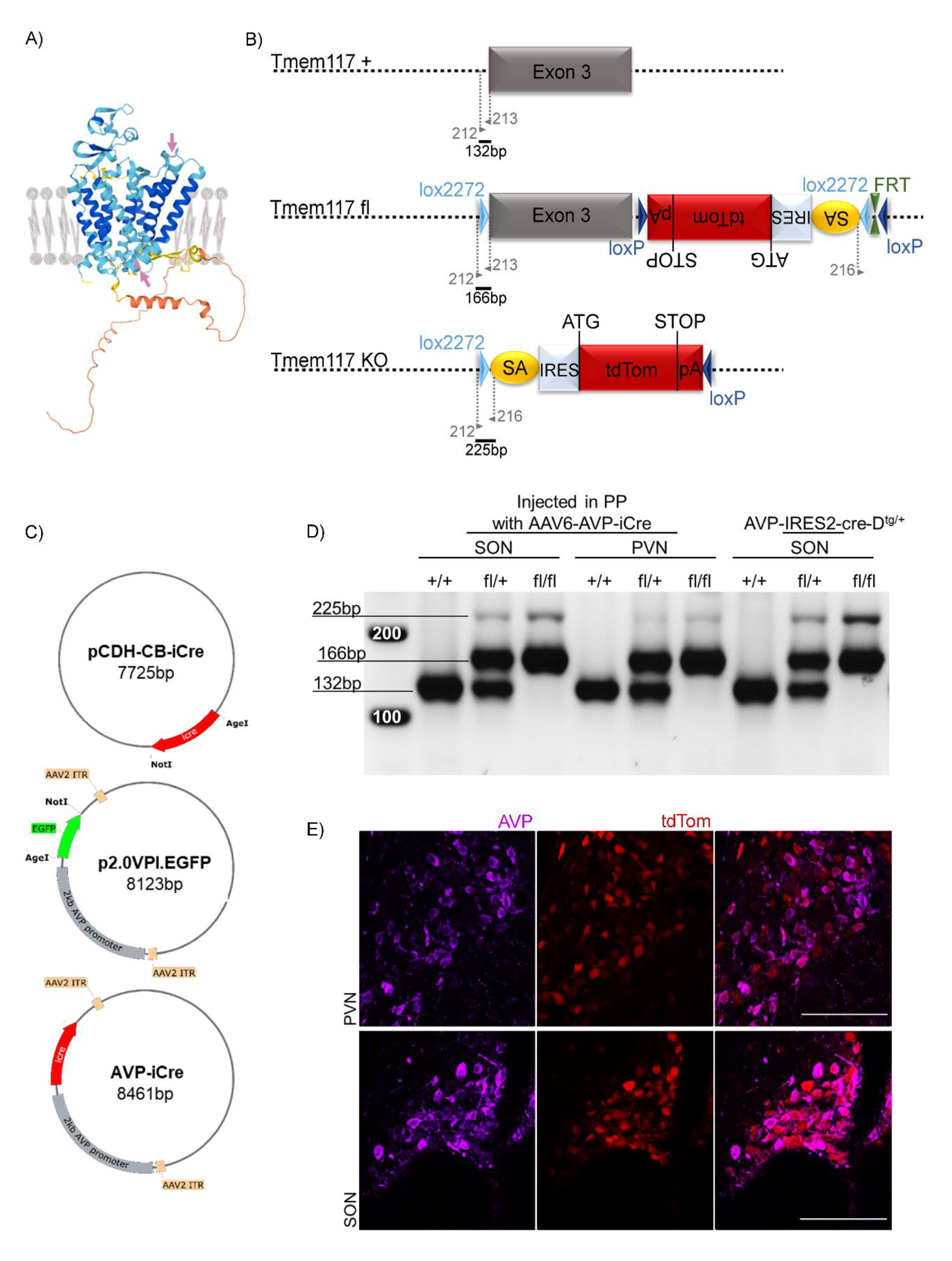
*Tmem117^fl^* mice and viral construct generation. (A) Predicted structure and transmembrane localization of the mouse Tmem117 (adapted from AlphaFold). The pink aminoacids highlighted by the pink arrows are the first (aa 93) and the last (aa 136) encoded by the exon 3 sequence that corresponds to the whole 3^rd^ transmembrane helix. (B) Schematic representation of the *Tmem117* wild-type locus (+), the *Tmem117^fl^* locus (fl), and the recombined *Tmem117* locus (KO). Primer binding sites and expected amplicon sizes are indicated. (C) Generation of AAV-AVP-iCre plasmid. The pCDH-CB-iCre plasmid was used as a donor for the iCre sequence located between the NotI and AgeI restriction sites. The p2.0VPI.EGFP plasmid contained the full-length 2kb AVP promoter in an AAV vector. The GFP-containing NotI-AgeI fragment of this plasmid was replaced with the iCre sequence to generate the AAV-AVP-iCre plasmid. (D) PCR analysis of *Tmem117* recombination. DNA extracted from microdissected PVN and SON of mice injected in the posterior pituitary with the AAV6-AVP-iCre showed a 225 bp fragment indicative of *Tmem117^fl^* recombination. SON DNA from *Tmem117^fl^* mice crossed with *AVP-IRES2-cre-D^tg/+^* mice was used as positive control. (E) Injection of AAV6-AVP-iCre in the posterior pituitary of *Rosa26^tdTomato/+^* mice led to efficient expression of tdTomato in AVP magnocellular neurons of the PVN and SON. AVP: Vasopressin, PP: Posterior pituitary, PVN: Paraventricular nucleus, SON: Supraoptic nucleus. Scale bar=100μm.

To assess the role of *Tmem117* in GCG secretion, *Tmem117^fl/fl^* and *Tmem117^+/+^* mice were injected with the AAV6-AVP-icre in the posterior pituitary to generate AVP*^TM117KO^* and AVP*^TM117WT^* mice, respectively (Fig 2A). Two weeks later the mice received an intraperitoneal (i.p.) injection of saline and blood was collected one hour later for plasma CPP and GCG measurements. The same experiment was repeated one week later with injection of insulin instead of saline to induce hypoglycemia. Insulin induced the same hypoglycemic levels in AVP*^TM117WT^* and AVP*^TM117KO^* male mice (Fig 2B); CPP secretion was increased in both groups of mice, but significantly more in AVP*^TM117KO^* mice than in AVP*^TM117WT^* mice (Fig 2C); the same pattern was observed for the secretion of GCG (Fig 2D).

**Figure 2.**
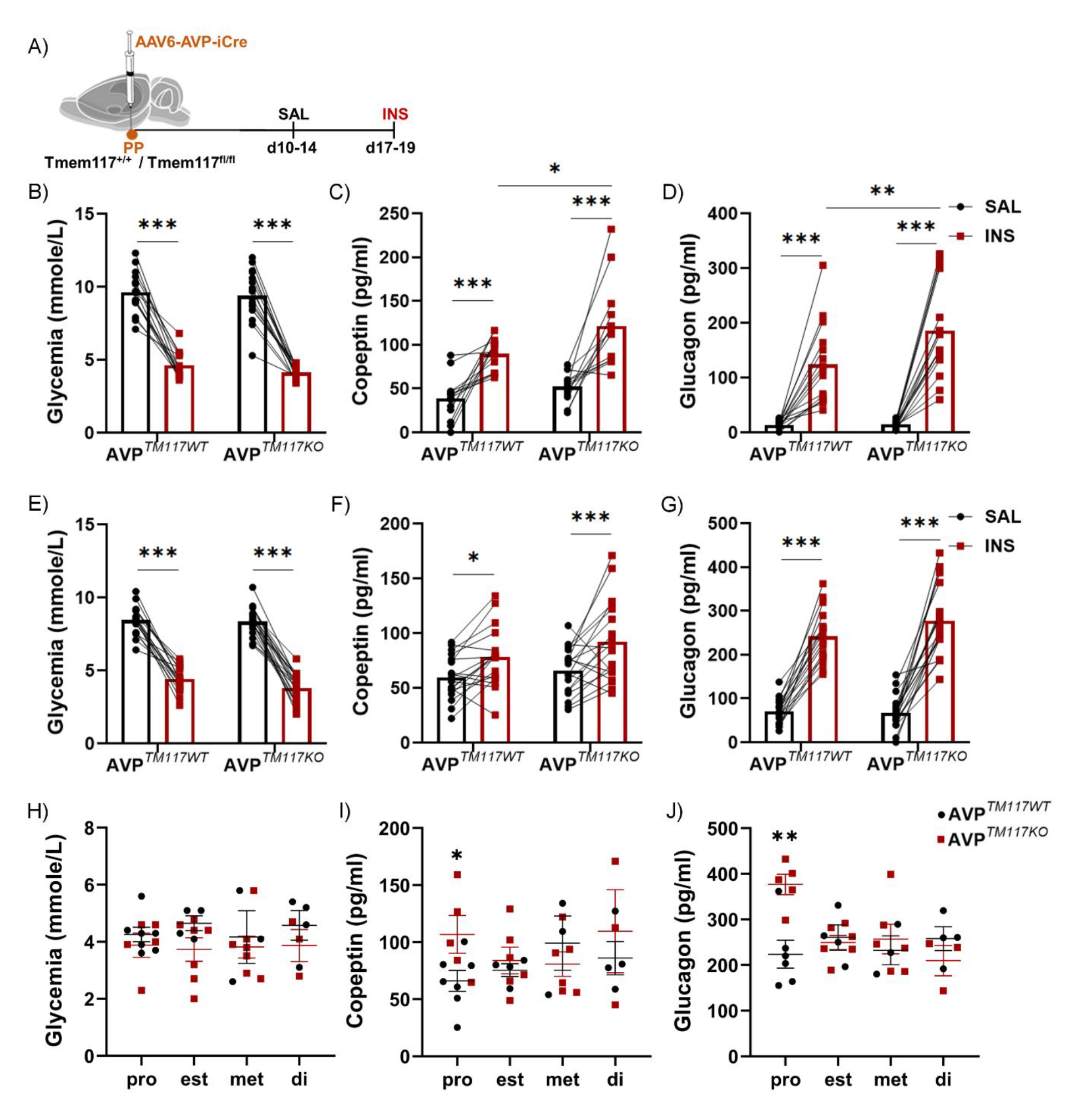
*Tmem117* inactivation in AVP neurons enhances hypoglycemia-induced copeptin and glucagon secretion. (A) Experimental scheme. AAV6-AVP-iCre was injected in the posterior pituitary of Tmem117^fl/fl^ (AVP^TM117KO^) or Tmem117^+/+^ (AVP^TM117WT^) mice. (B) Glycemia one hour after saline (black) or insulin (red) i.p. injection [n=15-16 mice per group]. (C) CPP plasma levels one hour after saline or insulin injection [n=13 mice per group; mean±SEM for INS: WT 89±5 vs KO 121±13 pg/ml]. (D) GCG plasma levels one hour after saline or insulin injection [n=15-16 mice per group; mean±SEM for INS: WT 124±19 vs KO 186±21 pg/ml]. (E) Glycemia one hour after saline (black) or insulin (red) i.p. injection [n=18-22 mice per group]. (F) CPP plasma levels one hour after saline or insulin injection [n=18-19 mice per group; mean±SEM for INS: WT 78±6 vs KO 92±8 pg/ml]. (G) GCG plasma levels one hour after saline or insulin injection [n=17-20 mice per group; mean±SEM for INS: WT 242±14 vs KO 277±18 pg/ml]. (H-J) Analysis of glycemic levels and of CPP and GCG secretion following insulin injection in female mice at each stage of the estrus cycle [n=3-7 mice per group]. (H) Glycemic levels (I) CPP plasma levels (J) GCG plasma levels. d: day, di: diestrus, est: estrus, INS: insulin, met: metestrus, PP: posterior pituitary, pro: proestrus, SAL: saline. For panels H-J, lines correspond to the mean value per group and error bars represent ± SEM. B- G: 2-way ANOVA RM with Bonferroni post-hoc test; H-J: unpaired t-test; *p<0.05, **p<0.01, ***p<0.001.

Female AVP*^TM117WT^*mice and *AVP^TM117KO^* mice were then tested in the same conditions. Insulin induced the same level of hypoglycemia (Fig 2E) and triggered comparable secretion of CPP and GCG in both groups of mice (Fig 2F,G). However, when the hormone levels were analyzed separately for each phase of the estrus cycle, we found that while the insulin-induced hypoglycemic levels were comparable across the estrus cycle between genotypes (Fig 2H), the level of secreted CPP and GCG were significantly higher in *AVP^TM117KO^* mice as compared to AVP*^TM117WT^*mice during the proestrus phase (Fig 2I,J).

### Glucose responsiveness of SON AVP neurons is not affected by *Tmem117* inactivation

To assess whether AVP neurons were activated by insulin-induced hypoglycemia C57Bl/6N male mice were injected with a saline solution or with insulin and their brains were collected two hours later. Immunofluorescence microscopy analysis revealed that hypoglycemia (<3.9 mmol/L) robustly increased c-Fos expression in both PVN and SON (Fig 3B,E) with a ∼3- fold increase in the number of c-Fos^+^ AVP neurons (Fig 3C,F) compared to the saline injected controls (Fig 3A,D). Furthermore, analysis of microdissected PVN and SON tissue one hour after insulin injection revealed downregulation of *Tmem117* mRNA in the SON (Fig 3G-I).

**Figure 3.**
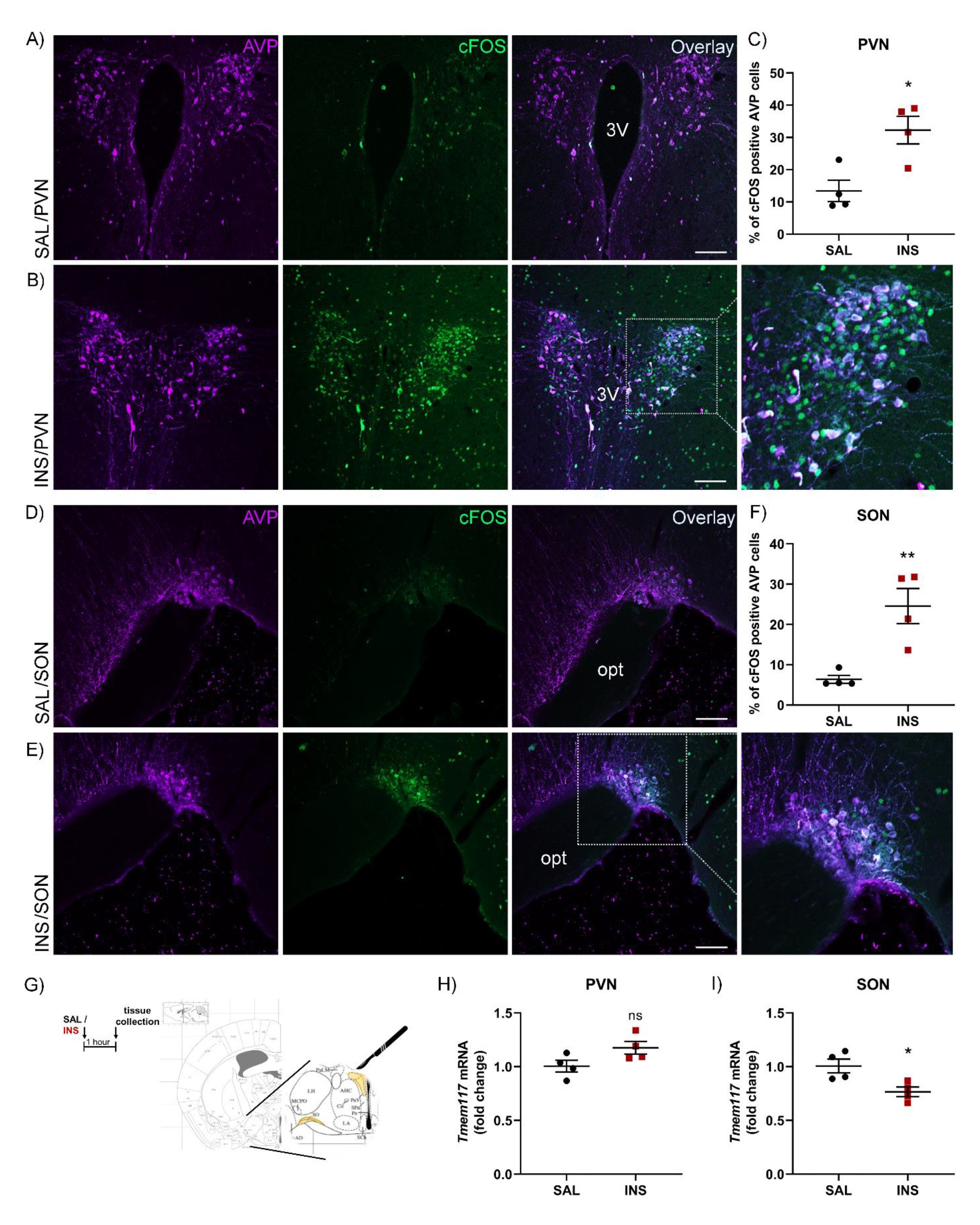
Insulin-induced hypoglycemia triggers activation of AVP magnocellular neurons and reduces *Tmem117* mRNA levels in the SON. (A-F) C57BL6/N male mice were injected i.p. with saline or insulin and c-Fos expression (green) in AVP (magenta) neurons was quantitated by immunofluorescence microscopy 2 hours later. (A) C-Fos expression in the PVN after saline injection. (B) C-Fos expression in the PVN after insulin injection. (C) Quantitation of c-Fos positive cells in the PVN after saline and insulin injections [n=4 mice per group]. (D) C-Fos expression in the SON after saline injection. (E) C-Fos expression in the SON after insulin injection. (F) Quantitation of c-Fos positive cells in the SON after saline and insulin injections [n=4 mice per group]. (G-I) C57Bl/6N male mice were injected i.p. with saline or insulin and brain tissue was collected 1 hour later. PVN and SON were microdissected for quantification of *Tmem117* mRNA levels by RT-PCR. (H) *Tmem117* mRNA levels in the PVN [n=4 mice per group]. (I)*Tmem117* mRNA levels in the SON [n=4 mice per group]. 3V: 3^rd^ ventricle, AVP: Vasopressin, INS: insulin, opt: optic tract, PVN: Paraventricular nucleus, SAL: saline, SON: Supraoptic nucleus. Scale bar=100μm. Lines correspond to the mean value per group and error bars represent ± SEM. Unpaired t-test; ns: p>0.05, *p<0.05, **p<0.01.

To determine whether *Tmem117* inactivation would modify insulin-induced c-Fos expression in AVP neurons, we prepared AVP*^TM117KO^* mice and AVP*^TM117WT^* control mice by injection of AAV6-AVP-iCre in the posterior pituitary of *Tmem117^fl/fl^* and *Tmem117^+/+^* male mice (Fig 4A). Two weeks later the mice were treated as described above for the C57Bl/6N. Induction of c-Fos expression in AVP neurons of the SON was comparable between the two mouse groups (Fig 4B,C).

**Figure 4.**
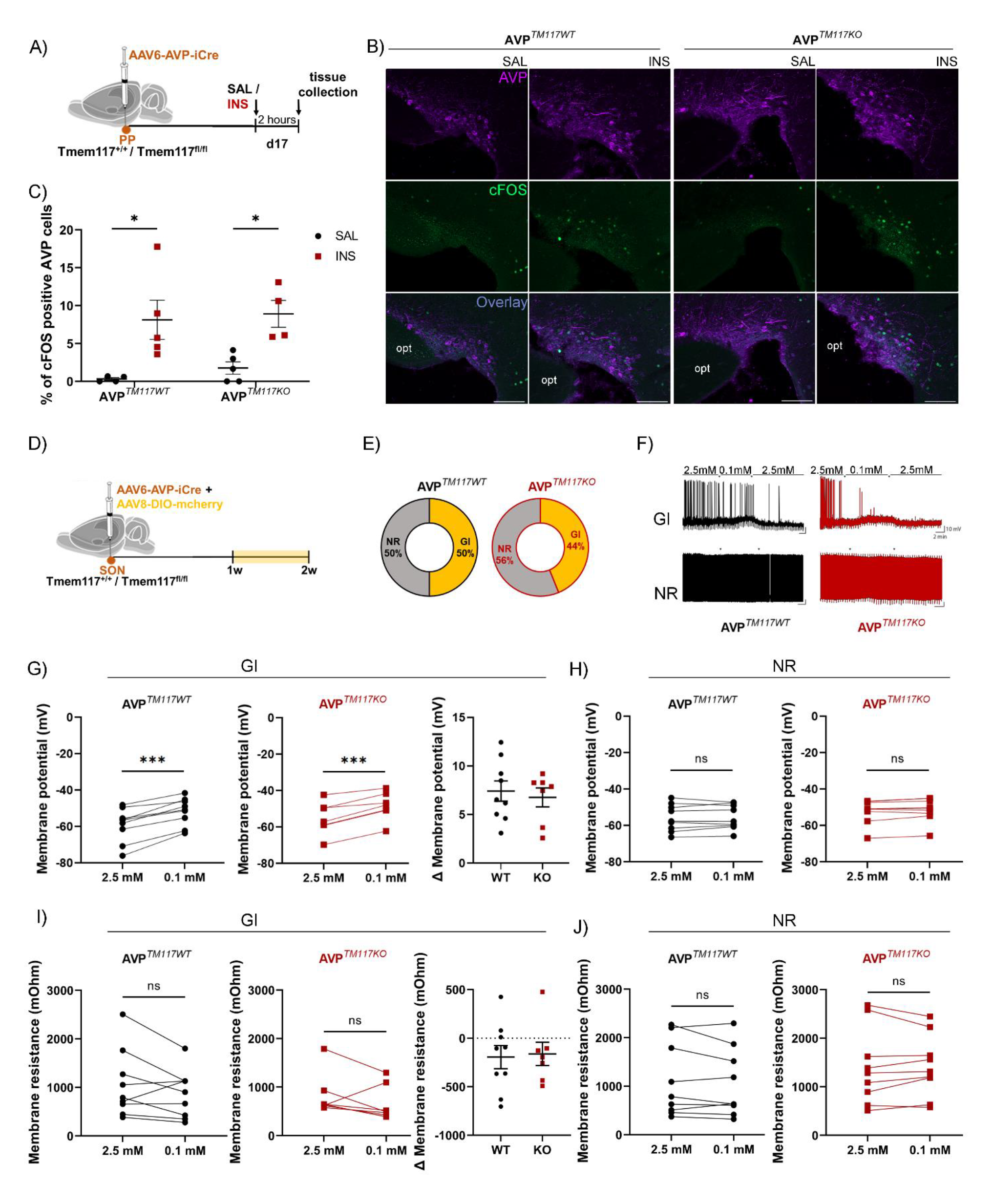
Glucose responsiveness of *Tmem117* KO AVP neurons. (A) Experimental scheme. AAV6-AVP-iCre was injected in the posterior pituitary of Tmem117^fl/fl^ (AVP^TM117KO^) or Tmem117^+/+^ (AVP^TM117WT^) mice. (B) C-Fos expression in the SON (C) Quantitation of c-Fos positive cells in the SON after saline and insulin injections [n=4-5 mice per group]. (D) Experimental scheme. AAV6-AVP-iCre was co-injected with AAV8-DIO-mcherry in the SON of Tmem117^fl/fl^ (AVP^TM117KO^) or Tmem117^+/+^ (AVP^TM117WT^) mice. (E) Proportion of GI and NR AVP neurons in the SON [n=16-18 cells per group]. (F) Representative traces of GI and NR AVP neurons in each group. (G) Membrane potential was increased comparably in response to 0.1 mM glucose in both groups. (H) No change in membrane potential in NR neurons of both genotypes [n=7-9 cells per group]. (I) Membrane resistance was decreased in 6 out of 9 AVP^TM117WT^ and 6 out of 7 AVP^TM117KO^ cells in response to 0.1 mM glucose. (H) Membrane resistance in NR neurons was stable for both genotypes [n=7-9 cells per group]. AVP: Vasopressin, d: day, GI: glucose inhibited, INS: insulin, NR: non-responding, opt: optic tract, SAL: saline, w: week. Scale bar=100μm. Lines correspond to the mean value per group and error bars represent ± SEM. C: 2-way ANOVA with Tukey post-hoc test; E: Fisher’s exact test; G-H: paired t-test for all graphs except Δ membrane potential (unpaired); ^ns^p>0.05, *p<0.05, ***p>0.001.

To determine whether AVP neurons were sensitive to the decreased glucose availability and whether *Tmem117* would modify this sensitivity, we performed patch clamp recordings of AVP neurons in the presence of 2.5 mM and 0.1 mM glucose (Labouèbe, Thorens and Lamy, 2018). *Tmem117^+/+^* and *Tmem117^fl/fl^* adult male mice were co-injected in the SON with AAV6- AVP-iCre and an AAV8-DIO-mCherry to allow fluorescence visualization of the AVP neurons (Fig 4D). SON was preferred over PVN due to its homogeneous magnocellular AVP population. Direct injection was used instead of retrograde labeling by injection in the posterior pituitary due to its higher labeling efficiency. One to two weeks after surgery, we performed patch clamp analysis of

AVP neurons. In brain slices of AVP*^TM117WT^* mice, we found that approximately half of the recorded AVP neurons were activated by low glucose (GI neurons) and that the other half were glucose non-responder (NR) neurons; no glucose excited (GE) neurons were observed (Fig 4E-H; black). When the same measurements were performed in cells with *Tmem117* gene inactivation, the proportion of GI and NR cells and their membrane potential responses were similar to that of AVP*^TM117WT^* mice (Fig 4E-H; red). Thus, AVP neurons were in large part GI neurons and *Tmem117* inactivation did not alter their glucose responsiveness. Furthermore, exposure of AVP neurons to low glucose concentration decreased the membrane resistance of 6 out of 7 *AVP^TM117KO^* GI cells and 6 out of 9 *AVP^TM117WT^* GI cells (Fig 4I) suggesting a cell autonomous response to hypoglycemia in those neurons.

### *Tmem117* inactivation increases ER stress, ROS production, intracellular Ca^++^ levels, and AVP mRNA expression

*Tmem117* has been identified in a siRNA screen of the HTC116 colon cancer cell line as a negative regulator of ER stress and ROS production (Tamaki *et al*., 2017). To verify its contribution to ER stress we used the colon adenocarcinoma cell line SW480, which endogenously expresses *Tmem117*. SW480 cells were transfected with a siRNA targeting *Tmem117* or a siRNA control and 48 hours later the cells were collected for RNA and protein extraction. A small reduction in *Tmem117* mRNA levels (log_2_fold ∼ −0.5) robustly increased expression of the ER stress markers *sXbp1* and *BiP* (Fig 3SA) and calnexin (Fig 3SB). *Tmem117* silencing in this cell line did not increase ROS production (Fig 3SC) but *Tmem117* overexpression significantly reduced ROS production (Fig 3SD). These data confirm a link between *Tmem117* expression and ER stress and ROS production in carcinoma cell lines.

**Figure S3.**
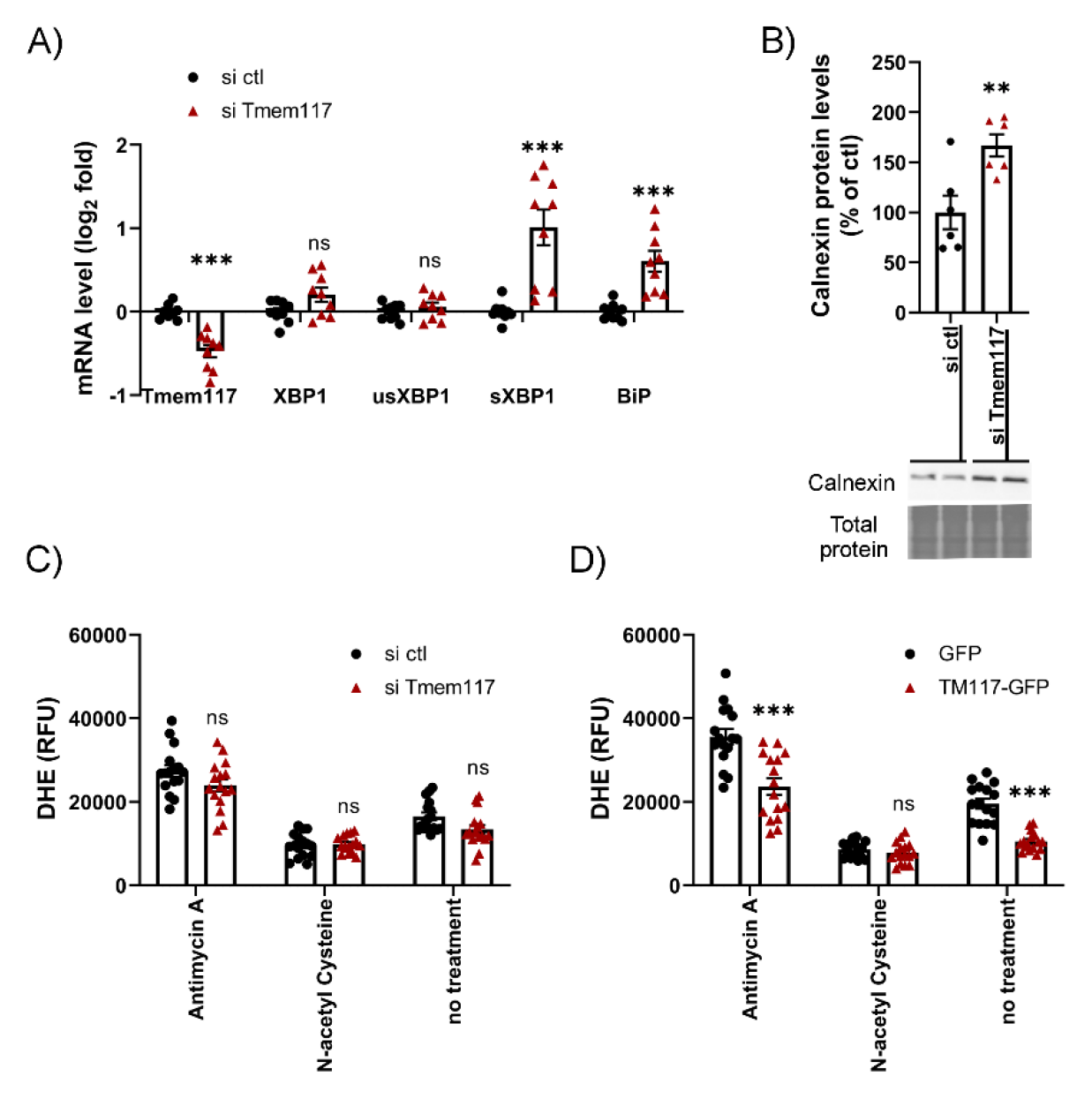
Tmem117 is modulating ER stress and ROS production in SW480 cells. (A) Expression levels of ER stress markers in SW480 cells. qRT-PCR quantification [n=8-9 samples/wells per group]. (B) Protein levels of calnexin in SW480 cells upon *Tmem117* silencing. Western blot quantification [[n=6 samples/wells per group]. (C) Intracellular ROS measurement in SW480 cells. DHE fluorescent signal in SW480 cells [n=16 samples/wells per group]. (D) Intracellular ROS measurement in SW480 cells upon overexpression of *Tmem117*. DHE fluorescent signal in SW480 cells [n=16 samples/wells per group]. DHE: dihydroethidium, RFU: relative fluorescent units. Bars correspond to the mean value per group and error bars represent ± SEM. Unpaired t-test; ns: p>0.05, **p<0.01, ***p>0.001.

To test whether inactivation of *Tmem117* in AVP magnocellular neurons *in vivo* would also affect ER stress and ROS production, we prepared *AVP^TM117KO^* and *AVP^TM117WT^* mice by injecting AAV6-AVP-iCre in the posterior pituitary of *Tmem117^fl/fl^* and *Tmem117^+/+^* mice. Two weeks later we collected their brains and measured the level of *Bip* mRNA by in situ hybridization (RNAScope). Quantitative analysis at the single-cell level revealed higher expression of *Bip* mRNA in AVP neurons of *AVP^TM117KO^* mice as compared to those of *AVP^TM117WT^* mice (Fig 5A,D); non-AVP cells in the SON showed no difference in *Bip* mRNA expression between the two groups of mice (Fig 5B,D). Furthermore, we found markedly elevated levels of AVP mRNA in AVP^TM117KO^ mice as compared to control mice (Fig 5C,D). AVP transcription has been reported to be induced by ROS (St-Louis et al., 2012, 2014). We, thus, assessed the ROS levels in the SON, by injecting i.p. the fluorescent ROS indicator dihydroethidium (DHE) in *AVP^TM117KO^* and *AVP^TM117WT^* mice twenty-four hours before tissue collection (Fig 5E). Quantification of the DHE fluorescence intensity revealed higher ROS production in the SON of *AVP^TM117KO^* mice (Fig 5F,G).

**Figure 5.**
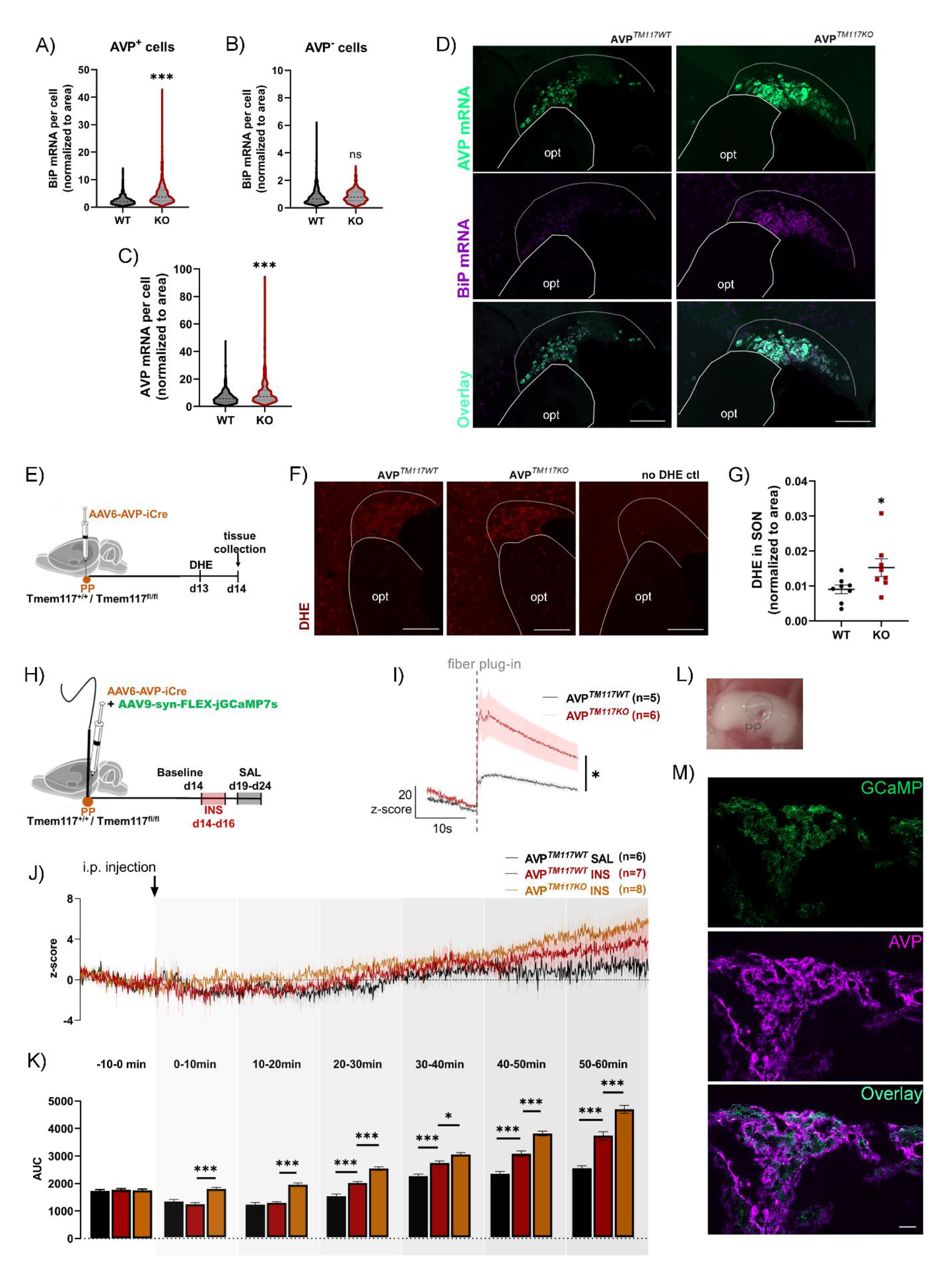
*Tmem117* inactivation in AVP neurons increases AVP and BiP mRNA, ROS levels and intracellular calcium in vivo. (A) Quantification of BiP mRNA per cell in AVP-positive cells of the SON [n=483-704 cells per group; each group consisted of 4 mice]. (B) Quantification of BiP mRNA per cell in AVP-negative cells of the SON [n=347-584 cells per group; each group consisted of 4 mice]. (C) Quantification of AVP mRNA per cell [n=483-704 cells per group; each group consisted of 4 mice]. (D) Fluorescence microscopy detection of AVP mRNA (green) and BiP mRNA (magenta) in the SON. (E) Experimental scheme. AAV6-AVP-iCre was injected in the posterior pituitary of Tmem117^fl/fl^ (AVP^TM117KO^) or Tmem117^+/+^ (AVP^TM117WT^) mice. DHE was injected i.p. (50mg/kg) 24 hours before tissue collection. (F) Fluorescence microscopy detection of DHE-derived fluorescence (red) in the SON. (G) Quantification of DHE-derived fluorescence in the SON [n=8 SON per group; each group consisted of 4 mice]. (H) Experimental scheme. AAV6-AVP-iCre was injected in the posterior pituitary of Tmem117^fl/fl^ (AVP^TM117KO^) or Tmem117^+/+^ (AVP^TM117WT^) mice. An optic fiber was implanted in the same area for signal analysis in freely moving mice. (I) Basal calcium levels measured in AVP magnocellular terminals of AVP^TM117WT^ and AVP^TM117KO^ mice. The gray dashed line represents the time at which the optic fiber cable was plugged onto the implanted cannula [n=5-6 mice per group]. (J) Calcium recordings in AVP magnocellular terminals 10 minutes before and one hour after the i.p injection of insulin (red and orange) or saline (black). The vertical arrow corresponds to the i.p. injection (time=0). (K) Quantification of mean signal intensity in 10-minute time bins. [n=6-8 mice per group]. (L) Stereoscope image of the pituitary gland depicting the optic fiber tract in the posterior pituitary. (M) Fluorescence microscopy detection of GCaMP (green) in posterior pituitary. Immunofluorescent labelling was used to localize the AVP terminals (magenta). AUC: area under the curve, AVP: Vasopressin, d: day, DHE: dihydroethidium, INS: insulin, opt: optic tract, PP: Posterior pituitary, SAL: saline, SON: Supraoptic nucleus,. Scale bar=20μm. For panels G and K, data are represented as mean ± SEM. For the violin plots of panels A-C, dashed lines correspond to the median value and dotted lines to the quartile values. A-C,G: unpaired t-test; I: 2-way ANOVA RM with Bonferroni post-hoc test; K: 1-way ANOVA with Tukey’s post-hoc test; *p<0.05, ***p>0.001.

Both ER stress and ROS can increase intracellular Ca^++^ concentrations (Giorgi *et al*., 2018), which would enhance neurosecretion. We, thus, measured the intracellular Ca^++^ levels in AVP neuronal terminals at the posterior pituitary, the site of AVP exocytosis, by *in vivo* fiber photometry. *AVP^TM117KO^* and *AVP^TM117WT^* mice were prepared by co-injection of the AAV6-AVP- iCre virus and an AAV9-hsyn-FLEX-jGCaMP7 in the posterior pituitary of *Tmem117^fl/fl^* and *Tmem117^+/+^* mice. During the same surgery an optic fiber cannula was implanted in the posterior pituitary for fluorescence monitoring (Fig 5H). Two weeks later the baseline jGCaMP7 fluorescence signal was recorded in both control and knock-out mice. This showed that inactivation of *Tmem117* induced a markedly higher baseline Ca^++^ signal (Fig 5I). We then performed Ca^++^ recordings after insulin-induced hypoglycemia. The first measures were performed at d14-16 after viral injection. A second session of recording was performed over d19- 24 in *AVP^TM117WT^* mice injected with saline and these data were used as control for the first set of experiments. For both sessions, the Ca^++^ signal was recorded during a 30-minute baseline period and for one hour following i.p. insulin or saline injections (Fig 5J). Each signal was normalized to the mean intensity of the baseline (corresponding to the last 10 minutes preceding the i.p. injection) and the area under the curve (AUC) was quantified for the mean trace of each group over 10-minute periods (Fig 5K). In *AVP^TM117WT^* mice insulin injection increased the Ca^++^ signal as compared to saline injection, a difference that became significant 20 minutes after the injection. In *AVP^TM117KO^* mice insulin induced a significantly higher Ca^++^ signal than in *AVP^TM117WT^* mice already 10 minutes after injection (Fig 5J,K). The correct placement of the optic fiber (Fig 5L) was assessed at the end of each experiment and the expression of jGCaMP7 in AVP magnocellular terminals was verified by fluorescence microscopy (Fig 5M).

Together, the above experiments indicate that *Tmem117* inactivation in AVP magnocellular neurons triggered ER stress, increased ROS production, intracellular Ca^++^ levels and AVP mRNA levels.

### *Tmem117* inactivation progressively triggers cell death

Permanently elevated ER stress and increased levels of intracellular ROS and Ca^++^ may trigger cell death (Zhivotovsky and Orrenius, 2011; Iurlaro and Muñoz-Pinedo, 2016; Giorgi *et al*., 2018). Therefore, we investigated the fate of AVP neurons one month after *Tmem117* inactivation in *Tmem117^fl/fl^* mice co-injected in the SON with AAV6-AVP-iCre and AAV9-hsyn-DIO-EGFP (*AVPT^M117KO^* mice). Two groups of control mice were prepared. One consisted of *Tmem117^+/+^* mice injected with both viruses (*AVP^TM117WT^* mice); these allowed to assess by immunofluorescence microscopy the total number of Tmem117^+^ cells and the total number of infected, EGFP-expressing cells. The second group consisted of *Tmem117^fl/fl^* mice injected only with AAV9-hsyn-DIO-EGFP (*AVP^TM117FL^* mice), which were used as control for the effect of the genotype on the total number of Tmem117^+^ cells in the SON (Fig S4A). Fluorescence microscopy images of EGFP and Tmem117 expression and of DAPI staining for the three groups are presented in Figure S4B.

**Figure S4.**
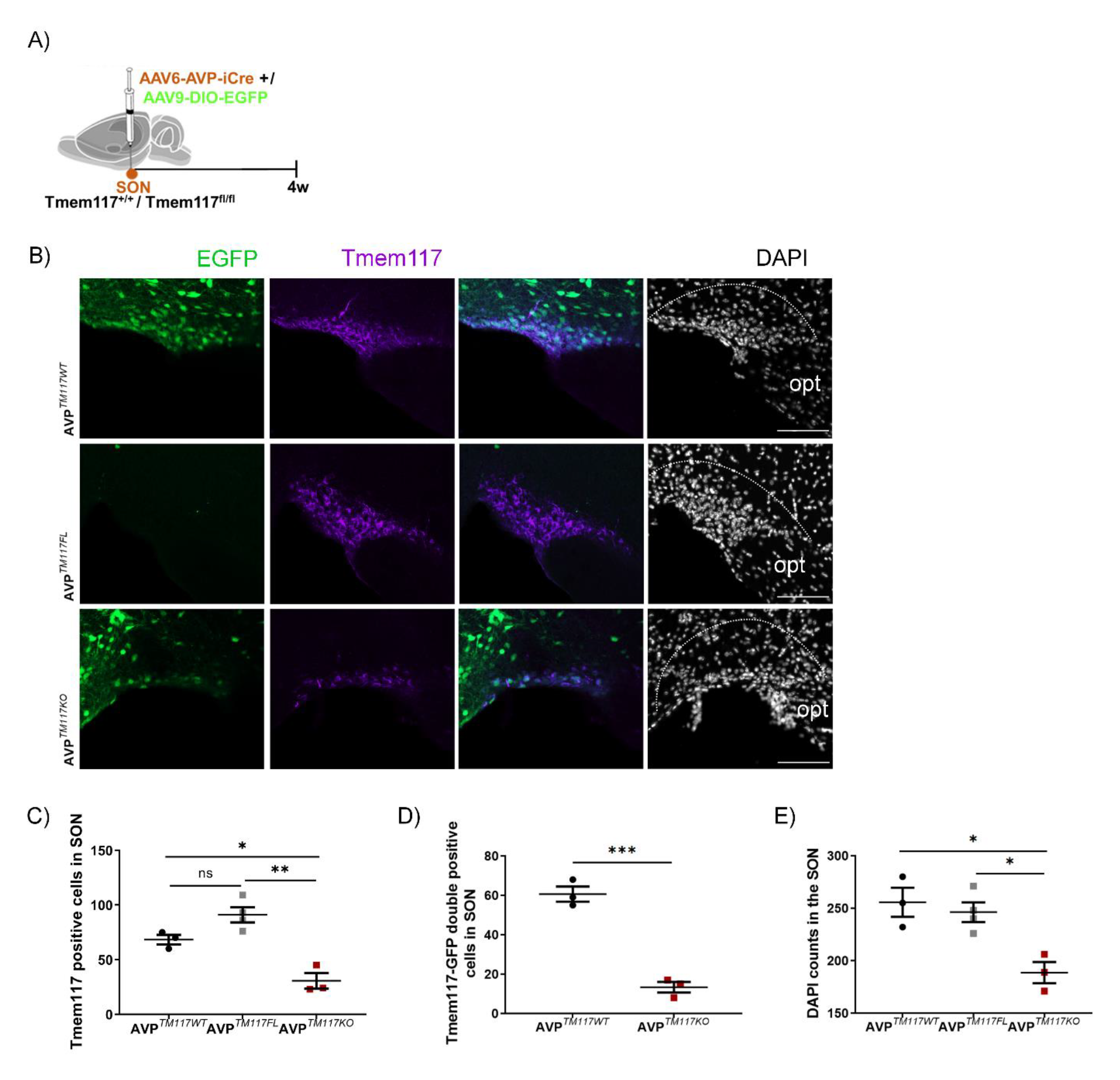
*Tmem117* inactivation in AVP neurons of the SON overtime leads to cell death. (A) Experimental scheme. AAV6-AVP-iCre was co-injected with AAV9-DIO-EGFP in the SON of Tmem117^fl/fl^ (AVP^TM117KO^) or Tmem117^+/+^ (AVP^TM117WT^) mice. AAV9-DIO-EGFP alone was injected in the SON of Tmem117^fl/fl^ (AVP^TM117FL^) to control for any genotype-derived effect. (B) Tmem117 (magenta), EGFP (green) and DAPI nuclear staining (white) in the SON. The dashed line is delimiting the SON area. (C) Quantification of Tmem117 expressing cells in the SON [n=3-4 mice per group]. (D) Quantitation of Tmem117 and EGFP double positive cells in the SON [n=3 mice per group]. (E) Quantitation of the number of cells (DAPI positive) in the SON [n=3-4 mice per group]. opt: optic tract, SON: Supraoptic nucleus, w: week. Scale bar=100μm. Data are represented as mean ± SEM. C,E: 1-way ANOVA with Tukey post-hoc test; D: unpaired t-test; *p<0.05, **p<0.01, ***p<0.001.

Quantitative analysis of the immunofluorescence data showed that the total number of Tmem117^+^ cells was the same in the two control groups but was reduced by ≥ 50% in *AVP^TM117KO^* mice as compared to the control mice (Fig S4C). When considering only the EGFP^+^ and Tmem117^+^ double labelled cells, the reduction was even more striking, reaching ∼75% (Fig S4D) indicating that most of the Cre-infected AVP neurons had disappeared during the post-infection period. This was also reflected by a reduction in the total number of neurons in the SON, as measured by DAPI staining (Fig S4E).

To determine whether the loss of AVP neurons would suppress the oversecretion phenotype, we injected AAV6-AVP-iCre in the posterior pituitary of *Tmem117^fl/fl^* (AVP*^TM117KO^*) and *Tmem117^+/+^* (AVP*^TM117WT^*) male mice and, except from the 2-week timepoint, we also assessed hormonal secretion upon insulin-induced hypoglycemia at the later timepoint of 1-month. Then, at the end of the experiment brains were collected for histological analysis (Fig 6A). Immunofluorescence microscopy analysis revealed a significant reduction in the number of AVP neurons in the PVN and SON of *AVP^TM117KO^* mice as compared to *AVP^TM117WT^* mice (Fig 6B-D). Insulin-induced hypoglycemia reached the same level in both groups (Fig 6E), and induced a transiently enhanced secretion of CPP and GCG in AVP*^TM117KO^* mice that was evident only at the early timepoint (Fig 6F,G). Correlation analysis between plasma CPP and GCG levels for each sample revealed a strong positive correlation (Fig 6H), further supporting their causal relationship. Thus, inactivation of *Tmem117* in AVP neurons led, over time, to cell death and the disappearance of the oversecretion phenotype.

**Figure 6.**
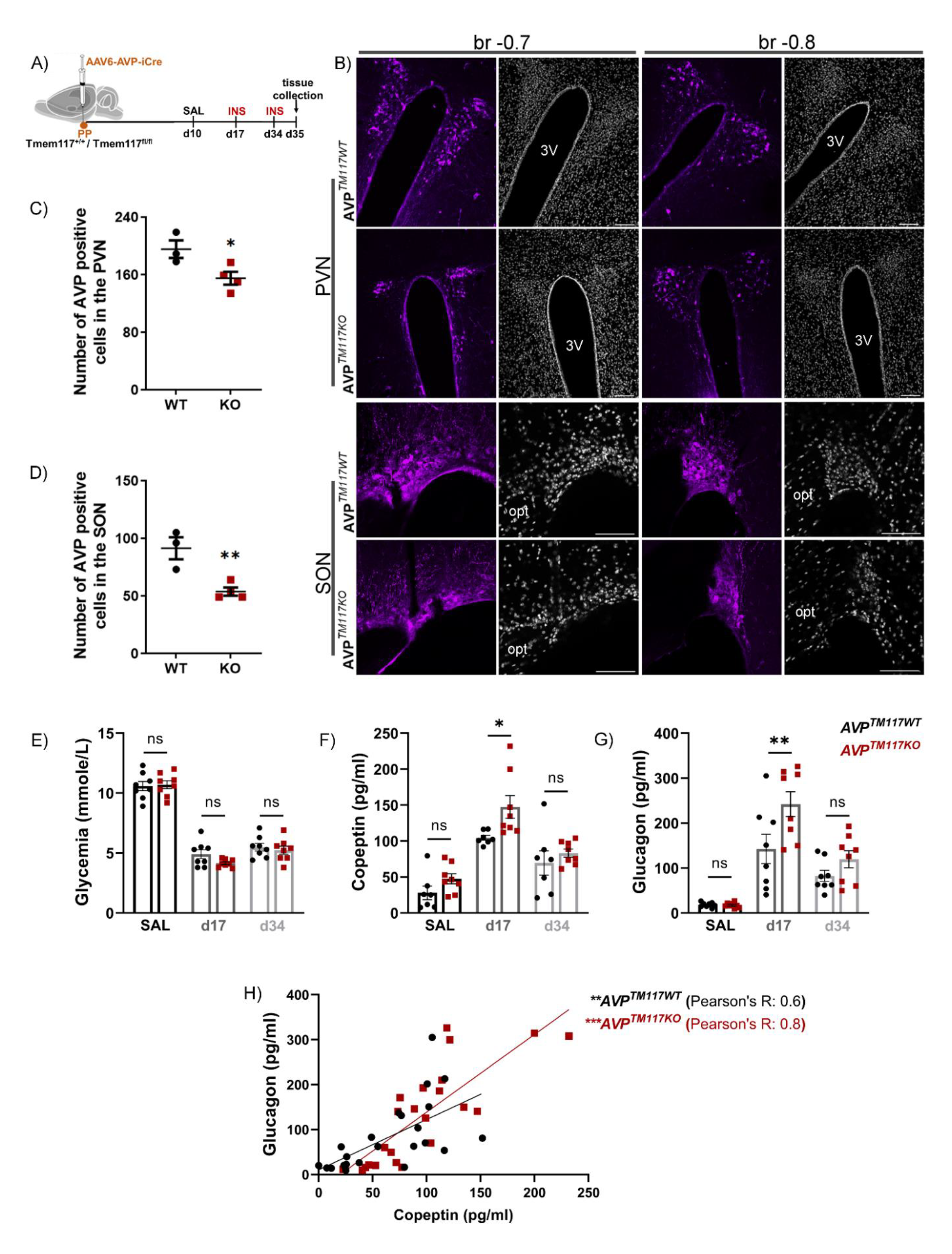
*Tmem117* inactivation in AVP neurons over time triggers neuronal death. (A) Experimental timeline. AAV6-AVP-iCre was injected in the posterior pituitary of Tmem117^fl/fl^ (AVP^TM117KO^) or Tmem117^+/+^ (AVP^TM117WT^) mice. (B) Immunofluorescence microscopy detection of AVP in the PVN and the SON. (C) Quantitation of AVP positive cells in the PVN [n=3-4 mice per group]. (D) Quantitation of AVP positive cells in the SON [n=3-4 mice per group]. (E) Glycemia one hour after saline (black) or insulin (red) i.p. injections in AVP^TM117WT^ and AVP^TM117KO^ male mice [n=8-9 mice per group]. (F) CPP plasma levels one hour after saline or insulin injection in AVP^TM117WT^ and AVP^TM117KO^ male mice [n=8 mice per group]. (G) GCG plasma levels one hour after saline or insulin injection in AVP^TM117WT^ and AVP^TM117KO^ male mice [n=8-9 mice per group]. (H) Correlation between the levels of plasma CPP and GCG in each sample [n=24 samples per group; 8 mice per group tested at 3 distinct timepoints]. 3V: 3^rd^ ventricle, AVP: Vasopressin, d: day, INS: insulin, KO: AVP^TM117KO^, SAL: saline, opt: optic tract, PVN: paraventricular nucleus, SON: Supraoptic nucleus, WT: AVP^TM117WT^. Scale bar=100μm. For panels C and D lines correspond to the mean value per group and error bars represent ± SEM. C,D: unpaired t-test; E-G: 2-way ANOVA for each timepoint in comparison to SAL baseline with Bonferroni post-hoc test; *p<0.05, **p<0.01, ***p<0.001.

## Discussion

In this study we characterized the site of expression and functional role of *Tmem117*, a genetically controlled, hypothalamic regulator of insulin-induced GCG secretion. We found that Tmem117 is expressed in AVP magnocellular neurons and *Tmem117* inactivation increases hypoglycemia-induced CPP and GCG secretion. This phenotype is observed in male mice and only in the proestrus phase in female mice. C-Fos immunodetection showed that AVP neurons are activated by hypoglycemia and patch clamp analysis revealed that about half of them are GI neurons. Inactivation of *Tmem117* did not affect the glucose responsiveness of AVP neurons, but instead increased ER stress, elevated intracellular ROS and Ca^++^ concentrations, leading to increased AVP production and secretion. These results highlight the physiological role of AVP neurons in the control of GCG secretion and identify *Tmem117* as a novel regulator of their function.

Tmem117 is an eight transmembrane domain-containing protein (Bürgi *et al*., 2016). We found it to be present in AVP neurons of the PVN and SON, with a widespread intracellular localization, from the soma to the axons and in their terminals located in the posterior pituitary, the site of AVP secretion in the bloodstream. The level of *Tmem117* mRNA expression in the hypothalamus of the BXD mouse lines used for the genetic screen showed strong and negative correlation with hypoglycemia-induced GCG secretion (Picard *et al*., 2022) (Fig S1D). Our physiological studies indicated that AVP neurons are activated by insulin-induced hypoglycemia, leading to CPP secretion, a response that was augmented by *Tmem117* inactivation and was accompanied by increased GCG secretion. These observations are in agreement with previous studies showing increased secretion of AVP in response to hypoglycemia (Baylis and Robertson, 1980; Baylis, Zerbe and Robertson, 1981; Chiodera *et al*., 1992) and that AVP stimulates GCG secretion through its binding to the AVP V1b receptor of pancreatic α cells (Dunning, Moltz and Fawcett, 1984; Spruce *et al*., 1985; Gao, Gérard and Henquin, 1992; Yibchok-anun *et al*., 2004; Kim *et al*., 2021; Liu *et al*., 2021). On the other hand, these results are also in line with our genetic screen that showed a negative correlation between *Tmem117* expression and insulin-induced GCG secretion.

Interestingly, the effect of *Tmem117* inactivation on GCG secretion is sex-dependent; it is present in male mice, but only during the proestrus phase in female mice. Circulating levels of estradiol peak during proestrus and have been linked to adaptations in AVP magnocellular neurons and their afferent connections (Sladek and Somponpun, 2008), as well as to decreased GCG secretion from pancreatic alpha cells (Godsland, 2005; Mårtensson *et al*., 2009). Thus, our data suggest a possible contribution of estradiol to the AVP-stimulated GCG release in conditions of hypoglycemia. Preceding studies have also reported that the CRR displays sex-dimorphism (Steinbusch *et al*., 2016; Briski *et al*., 2017), although the precise mechanisms involved still need to be fully characterized. Given the complexity and redundancy of the CRR implicated pathways, this sex-dimorphism is most probably a result of sex hormones acting at various sites and affecting different aspects of the response. Our data suggest that the action of estrogen on magnocellular AVP neurons is one of these aspects. AVP neurons express both nuclear and plasma membrane estrogen receptors that exert direct effects on intracellular signaling and AVP release (Sladek and Somponpun, 2008). Plasma concentration of AVP is decreased upon oestradiol treatment in rodents (Peysner and Forsling, 1990) and during the follicular phase in women (Stachenfeld, 2008) suggesting a suppressing effect of estrogen on AVP secretion. Therefore, one could hypothesize that hypoglycemia-induced AVP secretion would be decreased during estradiol-rich phases of the estrus cycle. Our data appear in line with such a hypothesis, since plasma CPP levels during hypoglycemia tend to be lower in WT mice that are in the proestrus phase (Fig 2I, black dots, p=0.13), but further studies focusing specifically on AVP secretion across the estrus cycle and its effect on CRR are required in order to get a detailed understanding of this interplay. What is clearly pointed out by our results is that inactivation of *Tmem117* in AVP neurons of female mice leads to increased AVP secretion during the estradiol-rich proestrus phase.

In order to investigate the mechanisms by which *Tmem117* inactivation regulates CPP secretion in hypoglycemic conditions, we explored the glucose responsiveness of the AVP neurons. First, c-Fos immunofluorescence analysis showed that insulin-induced hypoglycemia led to a ∼3 fold increase in the activation of AVP neurons in the SON, suggesting that they are glucose responsive. This was confirmed by patch clamp analysis, which revealed that ∼50% of them are GI neurons activated by low extracellular glucose concentrations. Thus, AVP neurons can respond to decreased glucose levels. Interestingly, a recent study reported that hypoglycemia can also activate AVP neurons through afferent connections arising from GI neurons of the basolateral medulla (Kim *et al*., 2021). Thus, AVP neurons are part of a brainstem-hypothalamus neuronal circuit where hypoglycemia can be sensed by neurons located at multiple sites to activate the secretion of AVP leading to increased secretion of GCG.

Patch clamp and c-Fos immunofluorescence analysis revealed that inactivation of *Tmem117* does not affect the glucose responsiveness of AVP neurons. Instead, it induces ER stress and increases intracellular ROS and Ca^++^ levels, leading to enhanced AVP production and secretion. These results extend previous observations made in the HCT116 cancer cells showing that *Tmem117* silencing increases ER stress and ROS production (Tamaki *et al*., 2017) and are in line with the reports that increased intracellular ROS stimulate AVP transcription in magnocellular neurons (St-Louis *et al*., 2012, 2014). It is striking that a small reduction in *Tmem117* expression in the SW480 cell line (∼30%) led to a significantly increased expression of ER stress markers. This suggests that *Tmem117* expression level has an important regulatory role. This is compatible with the genetic screen that showed strong correlation across the BXD lines between the level of *Tmem117* expression and hypoglycemia-induced GCG secretion. Thus, fine tuning of *Tmem117* expression appears as a physiological mean of controlling AVP neuron secretory capacity. This hypothesis is further supported by our data reporting decreased *Tmem117* mRNA levels in the SON in response to insulin-induced hypoglycemia. On the other hand, such a fine-tuning mechanism could also be implicated in pathological conditions, where a maladaptive upregulation of *Tmem117* expression would lead to decreased AVP secretion. In contrast, genetic inactivation of *Tmem117*, which leads to permanent increase in ER stress, ROS production and elevated intracellular Ca^++^ levels, is not compatible with the long-term survival of these neurons. Indeed, one month after induction of *Tmem117* recombination we observe neuronal death, probably as a result of apoptosis, as reported in the HCT116 cells with silencing of *Tmem117* (Tamaki *et al*., 2017).

Collectively, the present study highlights the physiological role of the AVP neurons in the CRR and identifies *Tmem117* as a genetic determinant of AVP and GCG secretion. Inactivation of *Tmem117*, did not affect the glucose sensing properties of AVP GI neurons, but led to increased CPP secretion associated with increased ER stress, intracellular ROS and Ca^++^ levels, and increased AVP mRNA expression. Thus, *Tmem117* emerges as a so far uncharacterized regulator of AVP neuron secretory capacity, by increasing AVP mRNA expression and secretion in response to hypoglycemia. Defining how Tmem117 controls these intracellular processes at the molecular level will, however, require future studies. Finally, this study further demonstrates that the central response to hypoglycemia is highly complex, integrating not only brainstem and hypothalamic glucose responsive neurons that activate the HPA axis and both branches of the autonomic nervous system, but also the secretion in the blood of AVP by magnocellular neurons.

## Materials & Methods

## RESOURCE AVAILABILITY

### Lead contact

Further information and requests for resources and reagents should be directed to and will be fulfilled by the Lead Contact, Bernard Thorens.

### Materials availability

Free of charge for noncommercial purposes

### Data and code availability

Data reported in this paper will be shared by the lead contact upon request. This paper does not report original code.

Any additional information required to reanalyze the data reported in this paper is available from the lead contact upon request.

## EXPERIMENTAL MODELS AND SUBJECT DETAILS

### Mice

All animal care and experimental procedures were in accordance with the Swiss National Institutional Guidelines of Animal Experimentation (OExA; 455.163) with license-approval (VD3363, VD3674) issued by the Vétérinaire Cantonal (Vaud, Switzerland). Mice were housed up to 5 per cage in a temperature-controlled room with a 12hr light/dark cycle and ad libitum access to water and standard laboratory chow (Kliba Nafag). Eight-to ten-week-old mice were used for all the experiments.

#### Generation of Tmem117^fl^ mice

Tmem117^fl^ mice were generated by homologous recombination in embryonic stem cells (genOway, Lyon, France). A reporter cassette for targeting exon 3, preceded by a splice acceptor was inserted in anti-sense into the respective intron. Both the cassette and the targeted exon were flanked by loxP sites and mutated loxP sites to enable deletion monitoring using the FLEx approach (Schnütgen *et al*., 2003) (FigS1A). The derived mouse line was developed onto a C57BL/6N background.

#### Tmem117^fl^; AVP-IRES2-cre-D mice

Tmem117^fl/fl^; AVP-IRES2-cre-D^tg/+^ mice were generated by crossing the Tmem117^fl^ line with the AVP-IRES2-cre-D (Jackson Laboratory code: 023530) for verifying the specificity of the Tmem117 antibody (Fig 1C,D).

#### Rosa26^tdTomato^ (Ai14) mice

Ai14 mice (Jackson Laboratory code: 007914) were used for verifying the specificity of the newly generated AAV6-AVP-icre.

### Cell lines

#### SW480 cells

The human colorectal cancer cell line SW480 (ATCC CCL-228) was cultured in Leibovitz’s L-15 Medium, supplemented with 10% fetal bovine serum (FBS). Cells were incubated at 37^◦^C in a 0.3% CO2 atmosphere and passaged twice per week up to 30 passages.

## METHOD DETAILS

### In vivo

#### Stereotaxic surgery / Optic fiber cannula implantation

Mice were anesthetized with a ketamine and xylazine solution (100mg/kg ketamine, 5mg/kg xylazine injected i.p.) and were placed on a stereotaxic frame (Stoelting, Chicago, USA). A small incision was made in the skin to reveal the skull. A small opening, to allow the passage of a 33 gauge needle, was made and the viral solution was injected at a rate of 100nl/min. The stereotaxic coordinates (relative to the bregma) and total volume injected for each area were: posterior pituitary: AP: −3, ML: 0, DV: −5.8, 0^◦^ angle / 400nl; SON: AP: −0.46, ML: ±1.15, DV: −5.45, 0^◦^ angle / 140nl per hemisphere. After the viral injection, the incision was closed by suturing and an analgesic solution was provided (0.1mg/kg buprenorphine s.c.). Ophthalmic ointment was used throughout the procedure to prevent eye dryness. A heat pad was used to maintain the proper body temperature.

For fiber photometry recordings an optic fiber (Doric Lenses B280-2408-6.4) was implanted in the posterior pituitary after the viral injection and the incision was closed with dental cement.

For insulin-induced hypoglycemia experiments 400nl of AA6-AVP-icre [8.1×10^12^ (viral genomes) vg/ml] were injected in the posterior pituitary. For electrophysiological recordings (Fig 4) and viability assessment (Fig S4) 140nl of a 1:1 mix of AAV6-AVP-icre (8.1×10^12^ vg/ml) with AAV8-syn-DIO-mCherry (2.2×10¹³ vg/ml) or AAV9-syn-DIO-EGFP (2.4×10¹³ vg/ml) were injected per hemisphere in SON. For fiber photometry 400nl of a 1:1 mix of AAV6-AVP-icre (8.1×10^12^ vg/ml) with AAV9-syn-FLEX-jGaMP7s (2.5×10¹³ vg/ml) were injected in the posterior pituitary.

#### Insulin-induced hypoglycemia; blood collection

Mice were food deprived for 6 hours (8am-2pm). At 12pm, they were placed in individual cages and glycemia was measured. At 1pm, glycemia was measured again and the mice were injected i.p. with either saline (baseline control) or insulin (0.8U/kg). At 2p.m. glycemia was measured again and 100-120μl of blood were collected under isoflurane-induced general anesthesia by submandibular vein incision.

#### Fiber photometry

Starting 5 days after surgery, mice were habituated to the recording cage and placement of the optic fiber in 30min sessions at least twice per week for two weeks. On the day of recording, the mice were placed in the recording cage for 30min and signal recording was started. The optic fiber was then connected to capture the baseline and stimulated signal intensity. The fiber photometry system was from Doric Lenses using LED light sources for GCaMP (465nm) and isosbestic (405nm) measurements; signal was recorded and analyzed using the Doric Neuroscience Studio software. The GCaMP7 signal was first normalized over the isosbestic signal (i=i_465_-i_405_) and then over the mean signal for the whole trace (ΔF/F0). Each point was then normalized to the baseline signal to obtain the corresponding z-score calculated using the formula: z_(i)_=[ΔF/F0_(i)_ – median(ΔF/F0 _baseline_)/ MAD_baseline_]. For baseline intensity traces z-score generation was calculated based on an 1 second recording period before the cable was plugged onto the optic cannula. For insulin-induced hypoglycemia traces z-score generation was calculated based on the baseline signal recorded over 10 minutes before the i.p. injection.

#### Estrus cycle monitoring

The estrus cycle phase was identified based on vaginal cytology samples after violet crystal staining (McLean *et al*., 2012).

#### Tissue collection

For immunofluorescence microscopy analysis mice were transcardially perfused with 10ml ice-cold phosphate buffered saline (PBS: 137mM NaCl, 2.7mM KCl, 10mM Na2HPO4, 1.8mM KH2PO4) followed by 40ml ice-cold paraformaldehyde (PFA, 4%) in PBS. Brain and pituitary were postfixed in 4% PFA O/N at 4^◦^C and then kept for 24 hours in 30% sucrose in PBS at 4^◦^C. Tissues were frozen and stored at −80^◦^C.

For SON and PVN DNA extraction, fresh brains were collected and kept in RNAlater solution (Thermo Fisher Scientific) at 4^◦^C for one week. Then, 250μm thick vibratome sections were prepared and PVN and SON nuclei were dissected under a stereomicroscope. Microdissected tissue was frozen with dry ice and stored at −80^◦^C, before DNA extraction.

### Ex vivo

#### Electrophysiological recordings

Tmem117^fl/fl^ and Tmem117^+/+^ mice were injected in the SON with an AAV6-AVP-iCre and an AAV8-DIO-mCherry. One to two weeks later, they were deeply anesthetized with isoflurane, decapitated, and their brain taken out and immediately placed in an ice-cold high glucose artificial cerebrospinal fluid (ACSF) containing (in mM): 125 NaCl, 2.5 KCl, 1.25 NaH2PO4, 1 MgCl2, 2 CaCl2, 26 NaHCO3, and 10 glucose (300 ± 5 mOsm) equilibrated with 95% O2 / 5% CO2. 250 μm coronal sections containing SON were prepared using a vibratome (VT1000S, Leica) and immediately transferred to an oxygenated ACSF solution containing 2.5mM glucose and maintained at 32°C for at least 1 h before starting the recordings. Experiments were performed using an upright epifluorescence microscope (BX51WI, Olympus, Japan) mounted on a motorized stage coupled to a micromanipulator (MPC-325, Sutter Instrument, USA) and equipped with a mercury lamp and an Evolve EMCCD camera (Teledyne Photometrics Technology, USA) and appropriate filters allowing the visualization of mCherry-expressing neurons.

Glucose responsiveness of AVP neurons was then assessed in the whole-cell configuration using a MultiClamp 700B amplifier associated with a 1440A Digidata digitizer (Molecular Devices). Borosilicate glass pipettes (resistance: 2-5 MΩ.; GC150F-7.5, Harvard Apparatus, USA) were prepared with a P-97 horizontal micropipette puller (Sutter Instrument, USA). The patch pipettes were filled with an intrapipette solution containing (in mM): 130 K-gluconate, 5 NaCl, 1 MgCl2, 10 NaPhosphocreatinine, 10 HEPES, 0.2 EGTA, 4 MgATP, 0.5 Na2GTP (pH 7.2-7.4; 280 ± 5 mOsm). Membrane potential and membrane resistance were monitored in current-clamp mode in the presence of 2.5 mM or 0.1 mM glucose after a 10–15 min baseline. Neurons with an access resistance >25 MΩ or changing by >20% during the recording were excluded from the analysis. Signals were filtered at 2 kHz, digitized at 10 kHz, and collected online with a pClamp 10 data acquisition system (Molecular Devices).

### Cell culture

SW480 cells were plated in 6-well tissue culture plate at a density of 300.000 cells/well. After 48 hours cells were transfected with siRNA against *Tmem117* or siRNA control (30pmol/well) with lipofectamine RNAiMAX (Thermo Fisher Scientific, Cat# 13778100). 48 hours after transfection cells were trypsinized and pelleted by mild centrifugation (1200rpm for 3min). Supernatant was removed and cell pellet was used for RNA and protein extraction using QIAGEN RNeasy Mini kit.

### Sample preparation / analysis

#### Immunofluorescence microscopy

25 μm-thick tissue sections were washed with PBS for 5min at RT, incubated with blocking solution (2% normal goat serum + 0.3% Triton x-100 in PBS) for 1 hour at RT and then incubated O/N with primary antibodies (Guinea Pig anti-(Arg8)-Vasopressin, BMA BIOMEDICALS, Cat# T- 5048; Rabbit anti-Tmem117, Novus, Cat# NBP1-94078) in blocking solution at 4^◦^C. Next, sections were washed with PBS three times for 10min, incubated with secondary antibodies (Goat anti-Rabbit Alexa Fluor 488, Molecular Probes, Cat# A-11078; Goat anti-Guinea pig Cy5, Abcam, Cat# ab102372) in blocking solution for 2 hours at RT, washed again with PBS three times for 10min, incubated with DAPI (1:10000 in PBS) for 20min at RT, washed with PBS three times for 10min, allowed to dry and finally mounted using fluoromount (Sigma F4680).

For c-Fos immunodetection tissue sections were washed with PBS for 5min at RT, incubated with blocking solution (4% normal goat serum + 0.3% Triton x-100 in PBS) for 1 hour at RT and then incubated O/N with primary antibody for c-Fos (Rabbit anti-cFos, Cell Signaling, Cat# 2250) in blocking solution at RT. Next, sections were washed with PBS three times for 10min, incubated with secondary antibody in blocking solution for 3 hours at RT, washed again with PBS three times for 10min and incubated O/N with primary antibody for AVP in blocking solution at 4^◦^C. Next, sections were washed with PBS three times for 10min, incubated with secondary antibody in blocking solution for 3 hours at RT, washed again with PBS three times for 10min, incubated with DAPI (1:10000 in PBS) for 20min at RT, washed with PBS three times for 10min, allowed to dry and finally mounted using fluoromount.

For all secondary antibodies the final concentration in the working solution was 1:500. For primary antibodies the concentrations used were: anti-Tmem117 1:250, anti-AVP 1:250, anti-cFos 1:1000.

#### In-situ hybridization (RNAscope)

25 μm-thick tissue sections containing the SON were co-stained by in situ hybridization for AVP mRNA (Cat# 401391) and BiP mRNA (Cat# 438831-C3) using RNAscope probes and RNAscope Fluorescent Multiplex Detection Reagents (Advanced Cell Diagnostics, Newark, CA, USA) following manufacturer’s instructions.

#### DHE fluorescent labelling

Tissue sections were washed with PBS for 5min at RT, allowed to dry and mounted using fluoromount (Sigma F4680).

#### Quantitative microscopy analysis

Fluorescence images were acquired on a ZEISS Axio Imager.M2 microscope, equipped with ApoTome.2 and a Camera Axiocam 702 mono (Zeiss, Germany). Specific filter cubes were used for the visualization of green (Filter set 38 HE eGFP shift free (E) EX BP 470/40, BS FT 495, EM BP 525/50), red (Filter set 43 HE Cy 3 shift free (E) EX BP 550/25, BS FT 570, EM BP 605/70), blue (Filter set 49 DAPI shift free (E) EX G 365, BS FT 395, EM BP 445/50) fluorescence and far red (Filter set 50 Cy 5 shift free [E] EX BP 640/30, BS FT 660, EM BP 690/50). Different magnifications were selected using a Zeiss x20 objective (Objective Plan-Apochromat 20x/0.8 M27, FWD=0.55mm) and a x40 oil-immersion objective (Objective C Plan-Apochromat ×40/1.4 Oil DIC M27 [FWD = 0.13 mm]). For super-resolution microscopy the ZEISS ELYRA7 SIM^2^ system was used with an x63 oil-immersion objective.

Cell quantification in PVN and SON was performed on images derived from two consecutive hypothalamic sections for each mouse (bregma −0.7 and −0.8). Fluorescence intensity quantification was performed using the QuPath-0.3.2 software on images derived from two consecutive hypothalamic sections for each mouse (bregma −0.7 and −0.8). Neuroanatomical landmarks (3^rd^ ventricle, optic tract, fornix) and DAPI staining were used to define the region of interest and cell area was marked manually by a blinded experimenter.

#### Western blot

The protein content of the samples was quantified by Bradford assay. 25μg of total protein per sample were prepared in SDS-PAGE sample buffer (2% SDS, 10% glycerol, 50mM Tris, 0.1% bromophenol blue, 5% β-mercaptoethanol) and separated on a 10% SDS-PAGE gel. Proteins were transferred onto a nitrocellulose membrane, incubated with blocking solution [3% milk in PBS-T (PBS + 0.1% Tween-20)] for 1 hour at RT, washed with PBS-T 3 times for 10min and incubated O/N at 4^◦^C with primary antibody (Rabbit anti-Calnexin, Abcam, Cat# ab133615) at a dilution of 1:1000 in PBS-T. Then membranes were washed with PBS-T 3 times for 10min, incubated for 1 hour at RT with horseradish peroxidase (HRP) conjugated secondary antibody (Anti-Rabbit HRP, Amersham, Cat# NA934) at a dilution of 1:10000 in blocking solution, washed again with PBS-T 3 times for 10min, incubated for 1min with enhanced chemiluminescence (ECL, Amersham) buffer and the HRP mediated chemiluminescence signal was detected/imaged using the Fusion FX6 Spectra imaging platform (VILBER). The derived images were analyzed for band intensity using imageJ. The calnexin signal of each sample was normalized to the total protein (Pierce Reversible Protein Stain Kit). Data are presented as percent of the control.

#### DNA extraction / PCR

DNA was extracted from microdissected tissue using the Arcturus Picopure DNA extraction kit (Applied biosystems, Ref # KIT0103). The extracted DNA was then subjected to PCR for detection of recombination using the following primers: 5’-CTTTCTTCATAAAAAGCCGGAA GGCATTAC-3’ (212), 5’-GCCTGAAATATAAATATCGCAAGTGAGTGTGC-3’ (213), 5’-CAACTGACCTTGGGCAAGAACATAAAGTG-3’ (216) (graphical representation Fig S1A).

#### RNA extraction / Real time-PCR

Total RNA was extracted from SW480 cell pellets using the QIAGEN RNeasy Mini kit based on manufacturer’s instructions and from microdissected tissue using the Arcturus Picopure RNA extraction kit (Applied biosystems, Ref # KIT0204). RNA was then reverse transcribed, and the derived cDNA was quantified using specific primers against *Tmem117* (FW:5’- TGTGATGCAGGACTGGGAAT-3’; RV:5’-TTGAACTGCATGTGAGGCGT-3’), *Xbp1* (FW:5’- TGGCCGGGTCTGCTGAGTCCG −3’; RV:5’-ATCCATGGGGAGATGTTCTGG −3’), *usXbp1* (FW:5’-CAGCACTCAGACTACGTGCA-3’; RV:5’- ATCCATGGGGAGATGTTCTGG-3’), *sXbp1* (FW:5’-CTGAGTCCGAATCAGGTGCAG-3’; RV:5’-ATCCATGGGGAGATGTTCTGG-3’), *BiP* (FW:5’-TGTTCAACCAATTATCAGCAAACTC-3’; RV:5’-TAGGTGGTCCCCAAGTCGAT-3’). Expression data were normalized to the housekeeping gene *Gusb* (FW: 5’- CCACCAGGGACCATCCAAT-3’; RV: 5’-AGTCAAAATATGTGTTCTGGACAAAGTAA-3’).

#### Quantification of circulating hormones

CPP and GCG levels in the plasma were quantified by Copeptin ELISA (MyBioSource, Cat# MBS2020621) and Glucagon ELISA (Mercodia, Cat# 10-1281-01), respectively, based on the manufacturer’s instructions.

### Constructs / Cloning / AAV packaging

The p2.0VPI.icre (AVP-icre) plasmid was generated by subcloning the iCre sequence (AgeI 2732nt, NotI 3813nt) from pCDH-CB-iCre plasmid (Addgene # 72257) in the p2.0VPI.EGFP backbone (AgeI 6237nt, NotI 6972nt; Addgene # 40868) (see Fig S1). The packaging of the plasmid in the AAV6 was performed by the University of North Carolina (Chapel Hill, NC) vector core.

The TM117-GFP plasmid was provided by the Van der Goot group (EPFL, Lausanne; Bürgi *et al*., 2016).

### eQTL analysis

eQTL mapping was performed using the R package R/qtl (Broman *et al*., 2003) with a genotype map from the BXD panel composed of GeneNetwork genotypes (www.genenetwork.org) merged with available RNA sequencing data from whole hypothalamic tissue (Picard *et al*., 2016). eQTL interval mapping was calculated using the expected-maximization algorithm, a 5% genotyping error rate, and pseudomarkers were generated every cM. eQTL location was obtained by 6.915 likelihood ratio statistics (LRS) support intervals. Significant eQTLs were determined for the trait using a 5% false discovery rate threshold estimated from 1000 permutations.

## QUANTIFICATION AND STATISTICAL ANALYSIS

All graphs and statistical analysis were generated using Prism software (GraphPad Prism 9), further details regarding sample size and the statistical analysis used in each case can be found in the corresponding figure legends. In brief, for experiments concerning comparisons between two groups on a single independent variable (Fig 2H-J; Fig 3C,F; Fig 4G^Δ^; Fig 5A,B,D-F,I; Fig 6C,DFig S3D,E) we used unpaired two-tailed t-test. For experiments concerning comparisons between two dependent measurements (Fig 4G,H) we used paired two-tailed t-test. For those concerning comparisons among three groups on a single independent variable (Fig 5M; Fig S3C,F) we used 1-way ANOVA with Tukey post-hoc test. For two-factor designs concerning repeated measures (Fig 2B-G; Fig 5K; Fig 6E-G) we used 2-way ANOVA RM with Bonferroni post-hoc test. For two-factor designs concerning non repeated measures (Fig 4C) we used 2- way ANOVA with Tukey post-hoc test. For the comparison of neuronal subpopulation proportion (Fig 4E) we used Fisher’s exact test. In all graphs error bars are depicting ±SEM.

## Acknowledgements

We thank Wanda Dolci for the excellent technical support. We thank Prof. G Van der Goot for kindly providing the TM117-GFP plasmid. We thank ZEISS and the EPFL Bioimaging and Optics Platform for the access to the ELYRA7 SIM^2^ super-resolution microscopy apparatus. This work was supported by a European Research Council Advanced Grant (INTEGRATE, No. 694798) and a Swiss National Science Foundation grant (310030-182496) to BT, and has received funding from the Innovative Medicines Initiative 2 Joint Undertaking (JU) under grant agreement No 777460 (HypoRESOLVE). The JU receives support from the European Union’s Horizon 2020 research and innovation program and EFPIA and T1D Exchange, JDRF, International Diabetes Federation (IDF), The Leona M. and Harry B. Helmsley Charitable Trust.

## Author Contributions

**SG:** conceptualization, methodology, investigation, formal analysis, visualization, project administration, writing-original draft, editing manuscript. **GL:** methodology, investigation, formal analysis, visualization, editing manuscript. **AP:** formal analysis, editing manuscript. **XB:** methodology, formal analysis. **ARSA:** software, visualization, editing manuscript. **BT:** conceptualization, supervision, project administration, resources, visualization, writing-original draft, editing manuscript, funding acquisition.

